# Novel Biomolecular techniques in the analysis of Orthopaedic trabecular metal implants produced by additive manufacturing

**DOI:** 10.1101/2025.02.12.637860

**Authors:** Jack Hurst, Richard OC Oreffo, Douglas Dunlop

**Affiliations:** Bone and Joint Research Group, Centre for Human Development, Stem Cells and Regeneration, Faculty of Medicine, University of Southampton, University Hospital Southampton NHS Foundation Trust

## Abstract

Recent trends have seen an increase in the use of orthopaedic acetabular implants produced by additive manufacturing.

These implants utilise novel trabecular metals with seemingly large variations in their structural designs.

There is no available biomolecular comparisons between these different implants. We have therefore created novel techniques to image and analyse 5 different implants.

Our results have shown differing responses of Human Bone Marrow Stem Cells cultured on these 5 different implants.

## Introduction

Advancements in 3D printing, known as additive manufacturing (AM), has allowed for the production of highly porous Trabecular Metals (TM) that aim to ‘mimic’ human bone allowing bony in-growth. AM is being increasingly used to produce acetabular implants with a TM surface to try and create a well-fixed component and reduce the rate of aseptic loosening.^1^

There is still much debate over the optimum structural design of these TMs, however, many ‘off the shelf’ implants already exist and are being widely used in arthroplasty surgery.^2^

From the limited data published on these products there is a wide variation in their structural properties, namely pore size, shape and porosity, as well as variations in the manufacturing process.

This highlighted the need for a study that can better evaluate, and potentially predict, the ‘real world’ performance of these developing materials to allow for accurate pre-clinical analysis, as well as direct comparisons between those implants already available.

## Materials and Methods

5 commonly available acetabular cups, all produced by AM were cut into multiple small samples. (Lima Delta TT, Adler Fixa-Ti-Por, Zimmer-Biomet G7 Osseo-Ti, Smith & Nephew Redapt, Stryker Trident II).

The samples were imaged with scanning electron microscopy (SEM) (FEI Quanta 250 SEM) to evaluate the trabecular structure and surface finish.

Human bone marrow stem cells (HBMSCs) were then seeded onto these samples, cultured for 21 days and then subsequently fixed for analysis.

Further cellular imaging was performed to visualise the stem cells on the trabecular surface, demonstrating the differences in attachment and proliferation across the different acetabular cups.

This was followed by biomolecular analysis to compare cell viability/proliferation (AlamarBlue Cell viability assay) and early markers of osteointergration (ALP Specific activity) across the different implants.

## Results

Biochemical analysis showed a statistically significant difference (P>0.05) in the early osteogenic response of HBMSCs cultured on the different implants. The Trident II sample showed a significant increase in specific ALP activity at all three timepoints, against all other samples, except for D7 against Redapt, which showed no significant increase (Fig.1).

**Figure 1:**
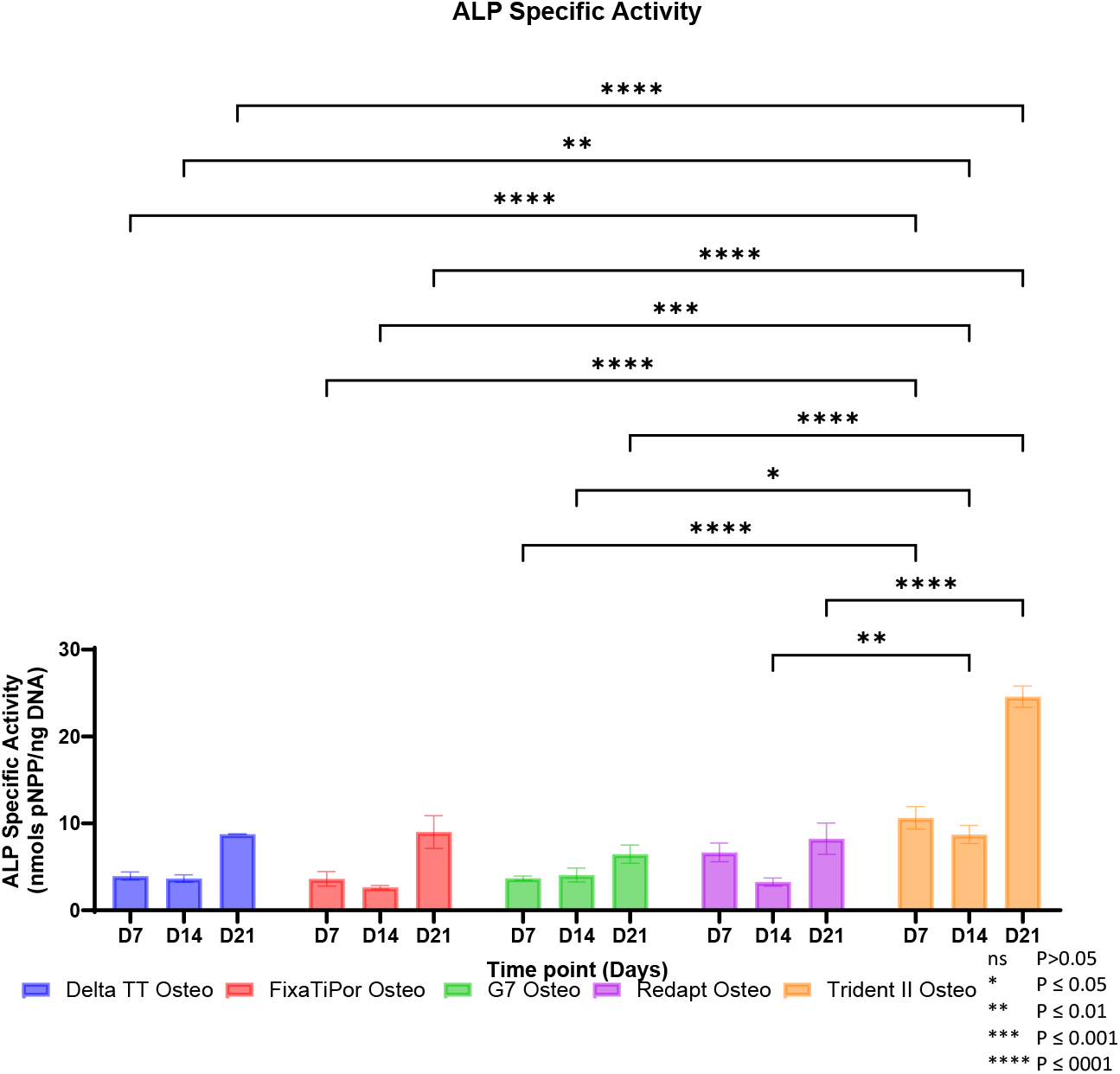
Graphical comparison of the Alkaline Phosphatase (ALP) Specific activity between the 5 different TM samples at 3 different time points (Days 7, 14 and 21).

The Trident II at day 11 showed a statistically significant increase in cell viability when compared to all samples except the Delta TT. The Delta TT also showed a statistically significant increase in cell viability when compared to the FixaTiPor, G7 and Redapt samples. The G7 showed a statistically significant increase only when compared to FixaTiPor samples. (Fig. 2).

**Figure 2.**
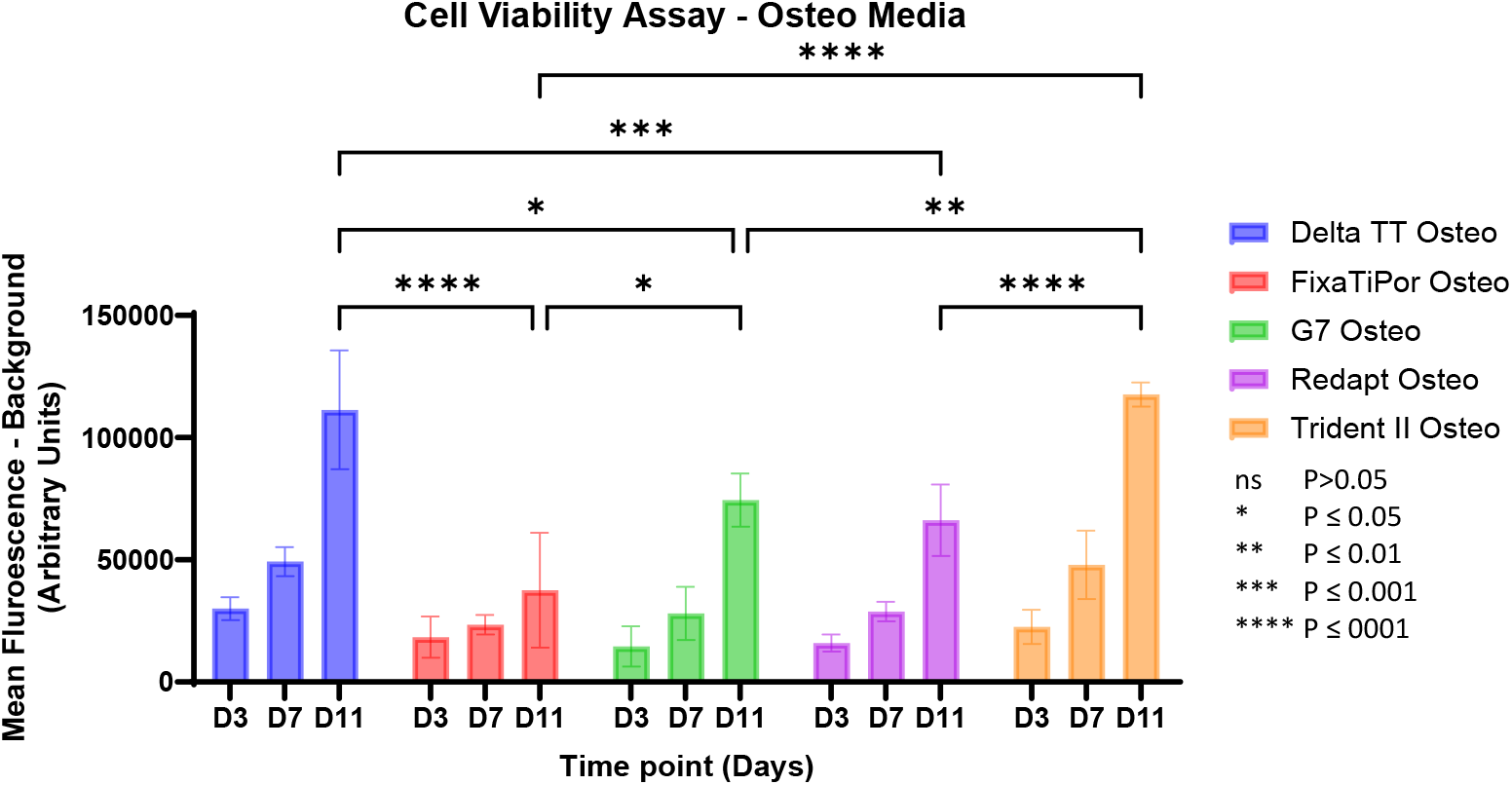
Graphical comparison of the Cell viability between the 5 different TM samples at 3 different time points (Days 3, 7 and 11) using and AlamarBlue Assay.

SEM images, shown in Figure 3, shows the wide variation in trabecular design across the different implants. At approximately 50x magnification the Delta TT and FixaTiPor have a very regular and repeating structure, compared to the much more random appearance of the G&, Redapt and Trident II implants. At a higher level of magnification (500x) the surface finish of each implant can be appreciated, with the FixaTiPor having a much smoother surface than that of the Trident II, G& and Redapt which all show a greater number of additional ‘beads’ adhered to the surface.

**Figure 3.**
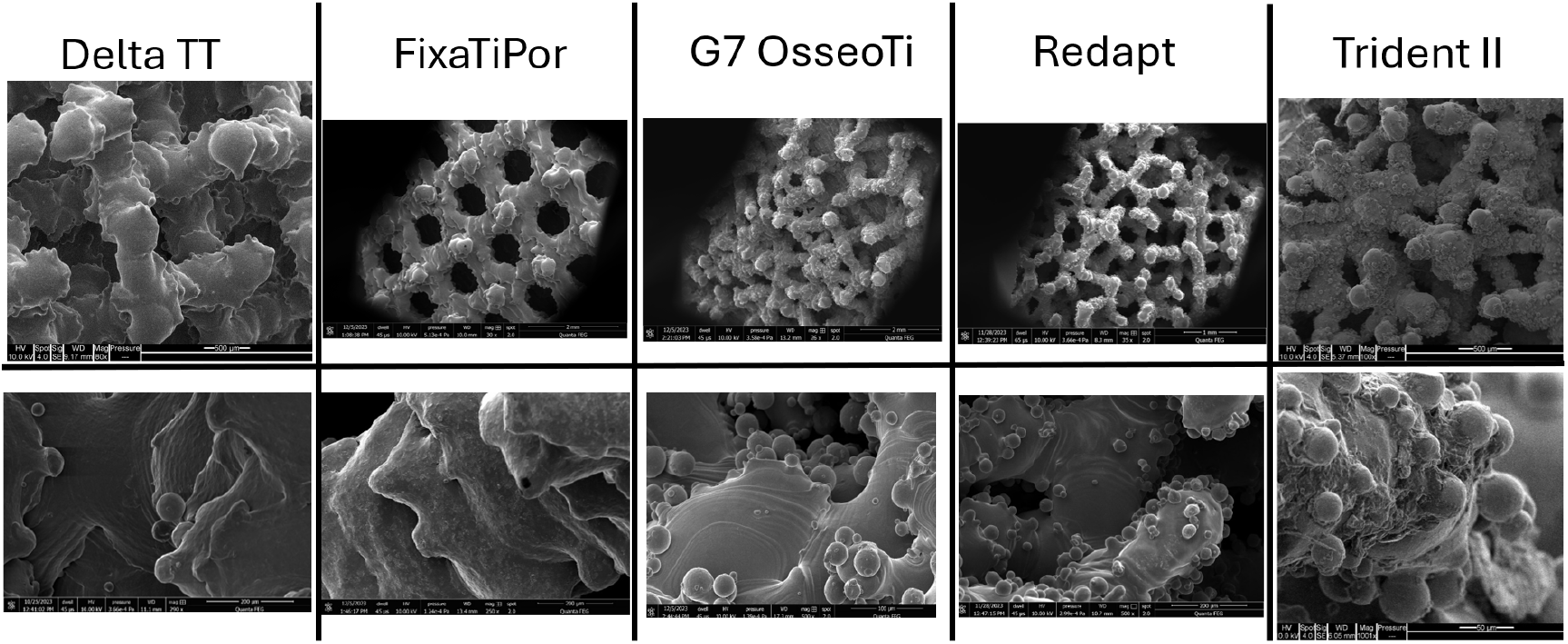
SEM images comparing the trabecular surface of 5 different TMs produced by AM. The top row is approximately 50x magnification and the bottom row approximately 500x.

Further SEM images, figure 4, performed following culture of the TMs with stem cells for 21 days shows how the cells attach across the surface. The difference in proliferation is also apparent with the Trident II showing a near complete ‘sheet-like’ covering of the surface compared to cells lining the pores of the TM as seen across the other samples.

**Figure 4.**
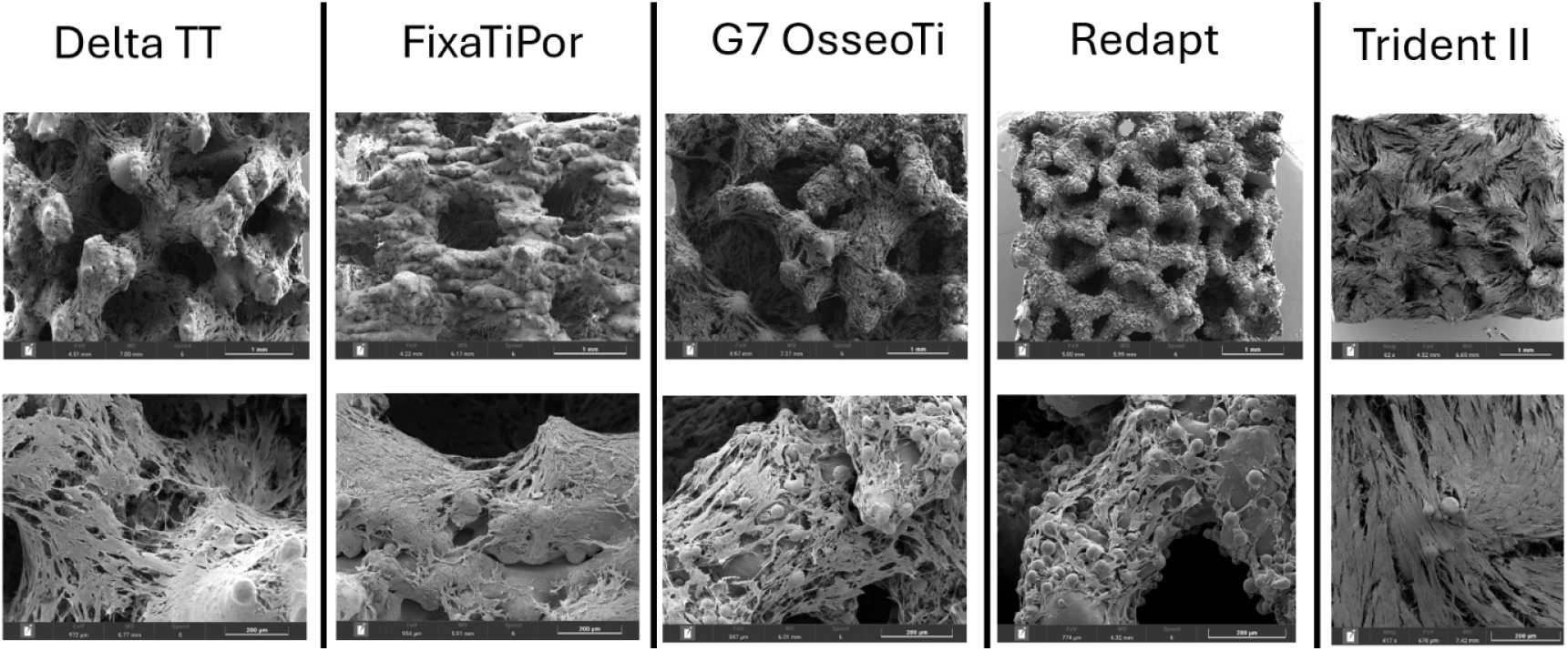
SEM images comparing the trabecular surface of 5 different TMs produced by AM after being seeded with HBMSCs and cultured for 21 days. The top row is approximately 50x magnification and the bottom row approximately 500x.

## Conclusions

These early results have shown a significant variance in the response of HBMSCs to each surface design/structure across the implants as highlighted by the biomolecular analysis.

The ALP specific activity analysis, an early marker of osteogenic differentiation on stem cells, shows the ability for stem cells to undergo osteogenic differentiation when cultured on these TM

The cell viability analysis not only shows the ability for the stem cells to proliferate over time on the different TMs it also shows significant differences between some of the different acetabular cups.

Whilst difficult to draw precise conclusions on the long-term performance of each implant this shows that the variation in TM design has an impact on HBMSC proliferation, differentiation and potential function.

This work is believed to be the first to perform biomolecular analysis and cellular imaging of HBMSCs on commercially available acetabular cups produced by additive manufacturing.

Future works will aim to correlate these varying structural designs with the differing cellular responses seen, ultimately to correlate these with long term clinical outcomes and performance.

## Notes

### Competing Interest Statement

The authors have declared no competing interest.

